# Genome-wide analysis and characterization of the LRR-RLK gene family provides insights into anthracnose resistance in common bean

**DOI:** 10.1101/2022.10.03.510363

**Authors:** Caroline Marcela da Silva Dambroz, Alexandre Hild Aono, Edson Mario de Andrade Silva, Welison Andrade Pereira

## Abstract

Anthracnose, caused by the hemibiotrophic fungus *Colletotrichum lindemuthianum*, is a damaging disease of common beans that can drastically reduce crop yield. The most effective strategy to manage anthracnose is the use of resistant cultivars. There are many resistance loci that have been identified, mapped and associated with markers in common bean chromosomes. The Leucine-rich repeat kinase receptor protein (LRR-RLK) family is a diverse group of transmembrane receptors, which potentially recognizes pathogen-associated molecular patterns and activates an immune response. In this study, we performed in silico analyses to identify, classify, and characterize common bean LRR-RLKs, also evaluating their expression profile in response to the infection by *C. lindemuthianum*. By analyzing the entire genome of *Phaseolus vulgaris*, we could identify and classify 230 LRR-RLKs into 15 different subfamilies. The analyses of gene structures, conserved domains and motifs suggest that LRR-RLKs from the same subfamily are consistent in their exon/intron organization and composition. LRR-RLK genes were found along the 11 chromosomes of the species, including regions of proximity with anthracnose resistance markers. By investigating the duplication events within the LRR-RLK family, we associated the importance of such a family with an expansion resulting from a strong stabilizing selection. Promoter analysis was also performed, highlighting cis elements associated with the plant response to biotic stress. With regard to the expression pattern of LRR-RLKs in response to the infection by *C. lindemuthianum*, we could point out several differentially expressed genes in this subfamily, which were associated to specific molecular patterns of LRR-RLKs. Our work provides a broad analysis of the LRR-RLK family in *P. vulgaris*, allowing an in-depth structural and functional characterization of genes and proteins of this family. From specific expression patterns related to anthracnose response, we could infer a direct participation of RLK-LRR genes in the mechanisms of resistance to anthracnose, highlighting important subfamilies for further investigations.

## Introduction

Common bean (*Phaseolus vulgaris* L.) has a significant importance in the current worldwide agricultural productivity scenario, especially in developing countries in South America, Central America, and southwest Africa^1^. Much of this importance is attributed to the versatility of this crop, which possesses nutritional, environmental, and economic benefits for producers and consumers^2^. However, due to its wide distribution and characteristic planting season, comprising several ecosystems and production systems, the bean crop is exposed to several factors that cause production instability, such as pests and diseases, which can be fungal, bacterial, or viral^2^,^3^.

Anthracnose is one of the most important diseases that affect the bean crop, caused by the hemibiotrophic fungus *Colletotrichum lindemuthianum*. This disease affects the quality of grains and pods, and can compromise up to 100% of the harvest in susceptible lines under favorable conditions for the pathogen development^1,4^. Genetic resistance is the most effective and safe way to control anthracnose in common beans^5^. Although more than 20 resistance loci, with independent effects, have been identified in different common bean linkage maps^5–8^, the genetic configuration for resistance in this pathosystem is challenging due to the great virulence diversity of *C. lindemuthianum*, which is represented by a large number of physiological races, which co-evolve with the host, allowing the selection of different gene combinations^4,9^. In this sense, several quantitative trait loci (QTLs) have also been mapped for resistance^10^.

Plant response to pathogen infections depends on the innate immunity of each cell and the systemic signs triggered from infection channels after the pathogen is recognized^11^. Thus, two layers of defense are used to activate the plant immune system. The initial mechanism uses transmembrane pattern recognition receptors (PRRs), which recognize and respond to molecular patterns associated with pathogen-associated molecular patterns (PAMPs) or microbe-associated molecular patterns (MAMPs). The reaction triggered by this first mechanism is known as PAMP triggered immunity (PTI). If the pathogen can overcome this initial response by not recognizing PRR receptors or by interrupting the signaling cascade and introducing its effector into the cell, the second response mechanism can be activated, which consists of effector-triggered immunity (ETI). Such a mechanism depends on the specific recognition of the pathogen’s effector by a host resistance R protein within the cell, leading to a hypersensitivity response^11^,^12^. R proteins play a key role at this second level of the plant immune system, both in pathogen recognition and signal transduction during the resistance response^13^.

Receptor-like protein kinases (RLKs) are a comprehensive superfamily of transmembrane receptors that sense stimuli on the cell surface and mediate cell signal transduction through autophosphorylation and subsequent downstream phosphorylation for intercellular communication^14^. The RLK superfamily is currently subdivided into 21 families, which stand out for their important role not only in the host defense system during plant-pathogen interactions but also during plant growth, development, and in response to abiotic and biotic stresses^15^,^16^.

Structurally, RLKs have a conserved serine/threonine kinase cytoplasmic domain (KD) and an amino-terminal extracellular variable domain (ECD). Most of the ECDs associated with RLK proteins have leucine-rich repeats (LRRs) that favor pathogen recognition due to their structural plasticity capable of detecting different ligands, e.g., proteins, peptides, and lipids^17^. One of the best-studied examples of PRRs in plants is Flagellin-sensitive 2 (FLS2), that recognizes the 22 conserved amino acids of bacterial flagellin in *Arabidopsis thaliana* (flg22) and forms a complex with its co-receptor Brassinosteroid-insensitive 1-associated receptor kinase 1 (BAK1) immediately after the perception to initiate the PTI response^18,19^. The participation of this protein was also verified, based on its expression pattern, in the defense system of *P. vulgaris* when interacting with the race 65 of *C. lindemuthianum*^20^.

LRR-RLKs constitute the largest family of the RLK superfamily and have been characterized in plants such as *A. thaliana, Oryza sativa, Brassica rapa, Solanum lycopersicum, Citrus* sp., and *Populus trichocarpa*^16,21–27^. However, despite the increasing characterization of LRR-RLKs in plants in recent years, to our knowledge, a comprehensive analysis has not yet been performed for *P. vulgaris*.

Characterizing a group of proteins in terms of their evolutionary history, structural and functional aspects, and associating this information with the analysis of their expression levels under specific conditions, enable a more accurate understanding of their participation in biological processes, such as plant-pathogen interaction and evolution of plants^27,28^. Furthermore, this information opens up possibilities for practical applications such as the pyramiding of different resistance alleles^12^ and the optimization of selection by identifying new loci and pathways related to disease resistance^29^. From this perspective, considering the increasing availability of genomic resources and biocomputational tools, this study aimed to characterize at a genomic scale the *P. vulgaris* LRR-RLK (PvLRR-RLKs) protein subfamily, as well as to study the dynamics of expression in the compatible and incompatible interaction of common bean plants with *C. lindemuthianum*.

## Material and Methods

### LRR-RLK Identification

The identification of the set of LRR-RLKs encoded by common bean was carried out based on the *P. vulgaris* kinome^30^. Initially, all putative protein sequences were downloaded from the *P. vulgaris* genome (v2.1) of Phytozome v13 (https://phytozome.jgi.doe.gov/pz/portal.html)^31^. The HMM profiles of typical kinase domains (PF00069 (Pkinase) and PF07714 (Pkinase_Tyr)) were downloaded from the Pfam protein family database^32^. For kinase protein (PK) identification, all protein sequences were aligned against these HMMs using HMMER v.3.3^33,34^ with an E-value cut-off of 0.1. A minimum coverage of 50% was established as the criterion for maintaining a sequence for further analysis. The identified PKs were subsequently classified into families and subfamilies based on the HMMs calculated with PK sequences of 25 plant species^35^. Only the major variant for genes with isoforms of the LRR-RLK subfamily proteins was maintained.

### Phylogenetic Analysis

To confirm the subfamily classification of the PvLRR-RLKs, we estimated a phylogenetic tree based on a multi-sequence alignment performed with the ClustalW software^36^. The tree was constructed using the evolutionary model of Tajima-Nei^37^ and the Neighbor-joining matrix-based reconstruction method considering 1,000 bootstrap replicates^38^. Both analyses were performed with the MEGA-X program (https://www.megasoftware.net) using the entire protein sequence and default parameters for each analysis^39^. The sequence of an LRR-RLK protein from *Chlamydomonas reinhardtii* (Cre06.g275450.t1.1), obtained from the phytozome, was used as an outgroup for phylogenetic analysis^40^, being identified in our analyses as Cr.LRR-RLK.

### Protein Properties

The biophysical properties of PvLRR-RLK proteins, including molecular weight (MW), isoelectric point (pI), and number of amino acids (aas), were calculated using the ExPasy ProtParam tool^41^. Additionally, to further verify the presence of LRR and KD domains, we employed the CDD tool (http://www.ncbi.nlm.nih.gov/Structure/cdd/wrpsb.cgi) ^42^, with default parameters and an E-value cut-off of 0.01. For the prediction and analysis of conserved motifs, the sequences were analyzed with the Multiple Em for Motif Elicitation - MEME Suite (version 5.3.3) (http://meme-suite.org/), considering a maximum motif number of 15 and an optimum motif width ranging from 6 to 50 amino acid residues^38,43^, with an E-value cut-off of 0.001. Visual inspections were performed with TBtools^44^.

The CELLO v.2.5 (http://cello.life.nctu.edu.tw/) ^45^ program was used to predict the subcellular location of PvLRR-RLK proteins. To illustrate the results, we created heatmaps in the TBtools^44^, highlighting the most likely location of each protein.

### Gene Organization and Chromosome Location

For the characterization of the gene organization, both the exon-intron organization and the analysis of regulatory cis-elements of the PvLRR-RLK genes were performed. The intron organization of all PvLRR-RLKs was evaluated using the Gene Structure Display Server - GSDS 2.0 (http://gsds.cbi.pku.edu.cn/) ^46^ considering the coding DNA sequences obtained from the Phytozome files. To evaluate the cis-elements associated with the promoters of the PvLRR-RLK genes, 1,500 base pairs (bps) upstream^47,48^ of the start codon were obtained from Phytozome and evaluated with the PlantCARE software (http://bioinformatics.psb.ugent.be/webtools/plantcare/html/) ^49^.

The distribution of LRR-RLKs across *P. vulgaris* chromosomes was assessed using the general feature format (GFF) file from the Phytozome database. Visual assessments of the PvLRR-RLK distribution across chromosomes were performed using the software PhenoGram (http://visualization.ritchielab.org/phenograms/plot) ^50^. Additionally, data from markers located 500 kb upstream and downstream to anthracnose resistance loci and QTL’s^51^ were associated with PvLRR-RLK gene positions and used to construct a chromosomal map using TBtools^44^.

### Duplication Events and Synteny Analysis

Duplication events and synonymous (Ks) and non-synonymous substitution (Ka) rates of LRR-RLKs across the *P. vulgaris* genome were estimated using the MCScanX toolkit implemented in TBtools^44^. The formula *T* = *Ks/*2*λ*, with *λ* representing the mean value of clock-like Ks rates (6.5 × 10^−9^) was used to calculate the date of duplication events^52^.

Additionally, we performed a synteny analysis of LRR-RLK genes between *P. vulgaris* and *Glycine max* Wm82.a4.v1 genomes, obtained from the Phytozome database. The syntenic blocks were estimated using the MCScanX toolkit, and the Dual Synteny Plot package was employed for visualization, both available in the TBtools program^44^.

### Gene Expression Analysis

The interaction between *P. vulgaris* and *C. lindemuthianum* was investigated regarding the expression levels of the LRR-RLK gene family. For this task, we employed transcriptomic data of common beans in the incompatible and compatible interaction with race 73 of *C. lindemuthianum*. The experiment was carried out for Padder et al.^53^, in a greenhouse with controlled temperature and humidity conditions. A pair of isogenic lines was used in the study, Jaguar possesses the Co-1 gene that provides resistance to race 73 of *C. lindemuthianum* whereas Puebla 152 lacks the resistance allele. The genotypes were inoculated with a conidia solution with a concentration of 2 × 10^6^ spores, and distilled water was sprayed on the control plants. Leaf samples were collected from both strains, inoculated and mock-inoculated, at 0, 24, 72, and 96 hpi at trifoliate growth stage. The RNA of the samples was extracted using TRIzol Kit (Invitrogen, Carlsbad, CA, USA), and mRNA-Sequencing (RNA-Seq) libraries using the Illumina TruSeq Stranded mRNA Library Preparation Kit following the manufacturer’s protocol. Sequencing was performed using Illumina HiSeq 2500 to generate single end (SE) reads of 50 nucleotides (nt)^53^.

These data are deposited in BioProject under code PRJNA342420^53^. From this library, data from 0 (control), 72, and 96 hours after inoculation (hpi) were used. The quality of the reads was evaluated with FASTqc^54^. The Kallisto tool^55^ was used to quantify gene expression. For this, the genome data of *P. vulgaris*, available at Phytozome (*P. vulgaris* v2.1), was used as a reference for the Kallisto index, followed by quantification for single-end libraries.

### Differential Expression Analysis of Anthracnose Resistance

After quantifying gene expression, we performed a differential expression gene (DEG) analysis using the DEseq2 R package version 1.18.1^56^. This analysis was performed by comparing data from resistant and susceptible plants at 72 and 96 hpi against data from the time/control condition at 0 hpi. For defining DEGs, we used a variance stabilizing transformation into read counts and performed a Wald test with a parametric fitting, considering a maximum p-value of 0.05 (FDR) and a minimum absolut log^2^ fold-change (based on the Transcripts Per Million - TPM metric) of 1.5. The minimum coverage of 1 (one) read was considered to calculate the saturation and expression. The fold-change calculations and p values (log2) were used to construct a volcano plot using the R statistical software^57, 58^. Venn diagrams were obtained using the Calculate and draw custom Venn diagrams tool (https://bioinformatics.psb.ugent.be/webtools/Venn/) to identify the intersection of DEGs between resistant and susceptible plants. Hierarchical cluster and heatmap analysis was performed in the R environment using the circlize R package with the k-means algorithm considering 5 clusters^59^.

## Results

### Identification of PvLRR-RLK genes

From the kinome of *P. vulgaris*^30^, 1,203 PKs were identified. Of these, only the proteins endowed with the transmembrane kinase and LRR domains were retained (Supplementary Table S1). All PvLRR-RLKs obtained were analyzed for redundancy following the criterion of maintaining the largest variants in the case of genes with isoforms. The nomenclature pattern obtained from Phytozome was maintained in subsequent analyses. After identification and initial screening, we considered a set of 230 PvLRR-RLKs for performing the analysis of this study. These proteins were classified into 15 subfamilies (from I to XV) according to the HMMER prediction, of which some subfamilies were further subdivided into other subgroups, as VI (LRR-VI-1 and LRR-VI-2), VII (LRR-VII-1, LRR-VII-2, and LRR-VII-3), X (LRR-Xa, LRR-Xb-1 and LRR-Xb-2), XI (LRR-XI-1 and LRR-XI-2), and XIII (LRR-XIII-a and LRR-XIII-b) (Supplementary Table S2).

Several characteristics of these proteins, including the number of aas, Mw, and pI, were also determined (Supplementary Table S3). There was a wide variation in the parameters evaluated, protein sizes ranged from 493 aas (Phvul.003G193100) to 1,290 aas (Phvul.006G198200), with a mean value of 855.56 aas (median of 882 and standard deviation of 181.355); the molecular weight ranged from 54,433.43 (Phvul.009G138532) to 140,774.43 (Phvul.007G067700), with a mean value of 73,307.67 (median of 72,847 and standard deviation of 797.84); moreover, the isoelectric point ranged from 5.11 (Phvul.004G173300) to 9.4 (Phvul.006G174700), with a mean value of 7.5 (median of 7.5 and standard deviation of 1.5).

### Phylogenetic analyses of PvLRR-RLKs

When analyzing the phylogenetic tree estimate, we observed that the classification of PvLRR-RLKs into subfamilies could be confirmed. Except for one isolated case (LRR-XI-1 subfamily), all proteins of the same subfamily were grouped together (Fig. 1). In this phylogenetic tree, the presence of an outgroup for the cr.LRR-RLK is clear, as expected. All the 230 LRR-RLKs analyzed were organized in a single group, which contained 23 subclades, representing the 22 different subfamilies. The LRR-XI-1 subfamily was the only one whose members were separated into two distinct subclades, one with 41 and the other with 17 proteins (Fig. 1). The other subclades, which grouped different subfamilies, showed varied sizes, with the smallest of them having only one protein (Phvul.009G171200) from subfamily LRR-XI-2. In contrast, the largest subclade had 40 and 41 proteins from the LRR-III and LRR-XI-1 subfamilies, respectively.

**Figure 1.**
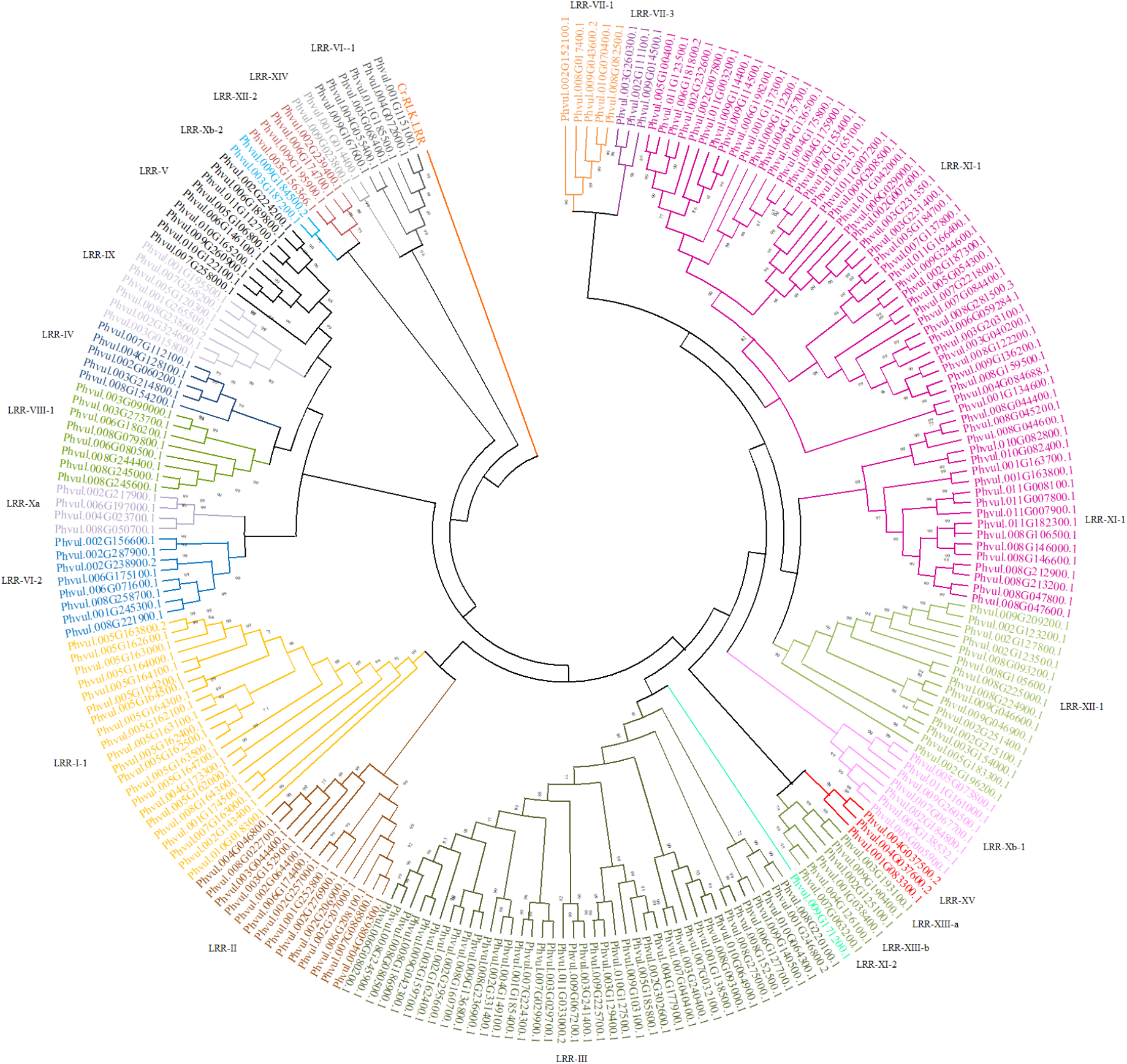
Phylogenetic analysis of LRR-RLK proteins of *Phaseolus vulgaris*. The different colors represent protein classification into subfamilies obtained by HMMER. Phytozome (https://phytozome.jgi.doe.gov/pz/portal.html) identification was maintained for all proteins. The outer group consists of an LRR-RLK protein from the alga *Chlamydomonas reinhardtii*, represented in the figure by Cr.LRR-RLK.

### Protein Properties

In order to gain insights into the diversity and functional characteristics of PvLRR-RLK proteins, the domain and conserved motif composition were analyzed using the CDD online tool and MEME, respectively, after which 16 conserved functional domains were identified (Supplementary Fig. S1). In addition to the typical kinase domains (Pkinase, and Pkinase_Tyr), the domains Pkc_like, LRRNT_2, and LRR_8 were the most common. As expected, at least one of these LRR and kinase domains was found in all proteins.

Interestingly, we found domains exclusively for specific PvLRR-RLKs, such as the Pkinase_fungal, TM_EphA1, and AsmA_2 domains, which were found in Phvul.002G206900 (LRR-II), Phvul.011G007800 (LRR-XI-1), and Phvul.008G045200 (LRR-XI-1) protein, respectively. In contrast, RIO1 was found in two proteins, Phvul.003G154000 (LRR-XII-1) and Phvul.009G184500 (LRR-Xb-2). In general, when we analyze the number and location of the domains in the proteins, we noticed that the domain pattern found was consistent between proteins of the same subfamily (Supplementary Fig. S1). For instance, in the LRR-I-1 subfamily, all proteins have, with exclusivity, the Malectin_like superfamily domain. Unlike this observation, some proteins differ structurally from others grouped in the same subfamily in one or more domains. In the LRR-XI-1 subfamily, protein Phvul.007G137800 was the only one with the APH domain. Observations like these can be made in all other subfamilies. As for characterization, the PK domains are larger. For example, the Pkinase domain had a size of 264 aas, whereas Pkinase_Tyr had 259 aas. In contrast, the LRR domains presented reduced sizes, e.g., the LRRNT_2 domain, with 41 aas, and the LRR8 domain, with 61 aas.

Motif analysis was performed with the MEME program to explore the evolutionary divergence of the KD and LRR domains. Fifteen motifs resulted from the analysis (Fig. 2). The members of the same subfamilies shared a similar motif composition in their domains (Supplementary Fig. S2). Motifs 1, 2, 8, and 14 were the most frequent, present in almost all proteins. Among these motifs, motif 1 belongs to the Pkinase domain, 2 to the Pkinase_Tyr, 14 to the LRRNT-2, and 8 to the LRR8 domain. However, the number and organization of these motifs vary among proteins grouped into different subfamilies. Variations in the presence of one motif were observed in some groups, such as in the LRR-XI-1, LRR-XII-1, and LRR-Xb-1 subfamilies, in which one or two proteins showed the motif 10 in their structure, whereas the other members of the same group did not present it.

**Figure 2.**
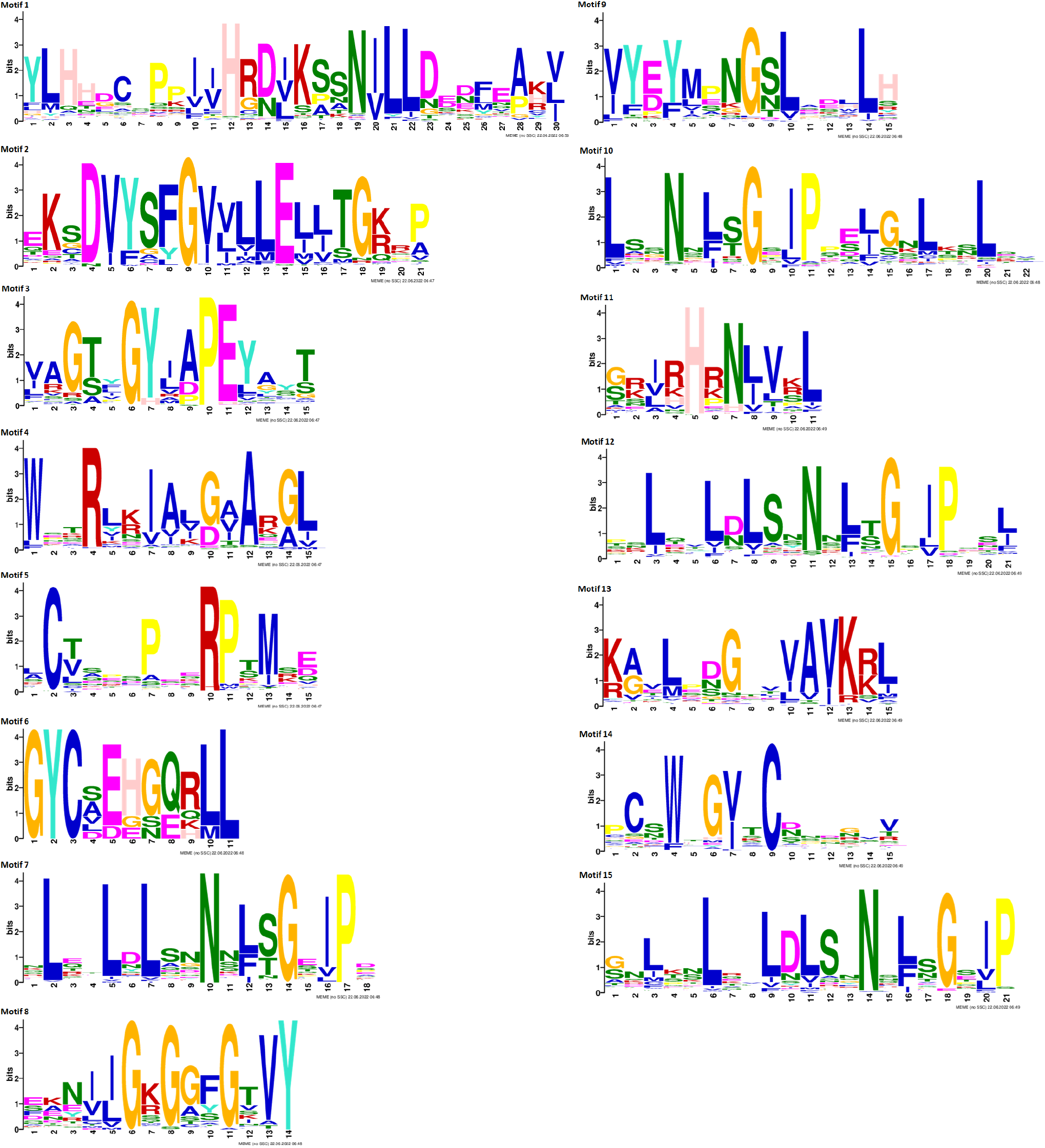
Conserved motifs, LRR domains, and consensus sequences of *Phaseolus vulgaris* LRR-RLK proteins. If the bit value of amino acid at this position is smaller than 1, it is represented with x; 2 *>* bits ≥ 1, with lowercase; 3 *>* bits ≥ 2, with capital letter; bits ≥ 3, with bold capital.

The subcellular location prediction obtained by CELLO v.2.5 showed that out of the 230 PvLRR-RLKs studied, 145 (63.05%) are located in the plasma membrane (Supplementary Table S3). Among the other proteins, the location was distributed in the extracellular space (46 proteins), cytoplasm (19), chloroplast (15), nucleus (7), and mitochondria (2) (Supplementary Fig. S3). Among the highlights, we can mention the case of the LRR-I-1, LRR-XIV, LRR-VII-3, and LRR-XIII-a subfamilies, which presented all the proteins predicted as having the localization in the plasma membrane. There was no consensus subcellular location for all representatives of the other subfamilies. In the subfamily LRR-VIII-1, the Phvul.009G043600 protein was the only one that differed in its subcellular location, being predicted in the chloroplast. In fact, most PvLRR-RLKs from each subfamily were predicted to act at the plasma membrane level. Some members, however, had their location predicted for other subcellular spaces.

### Gene Organization and Chromosome Location

The LRR-RLKs genes identified in *P. vulgaris* are distributed in its 11 chromosomes (Figure 3). In terms of absolute numbers, chromosome 08 (Chr 08) had the highest number of PvLRR-RLK genes, and Chr 10 the lowest (Table 1). The Phvul.L002151 gene, from the LRR-XI-1 subfamily, was located in a scaffold and, therefore, information about such a gene is not still available in some analyses (Supplementary Table S2, S3, S4, S7; Table 1). When analyzing the distribution of PvLRR-RLKs in the chromosomes, we observed the presence of these proteins in the telomeric regions, with emphasis on the concentration of proteins of the LRR-I-1 subfamily in the telomeric region of Chr 05, something not repeated at such intensity for other subfamilies and chromosomes. However, several examples of gene concentration in tandem of the same subfamily were recorded, as observed for subfamily LRR-XI-1 in the Chr 08, Chr 09, and Chr 11 (Fig. 3).

**Figure 3.**
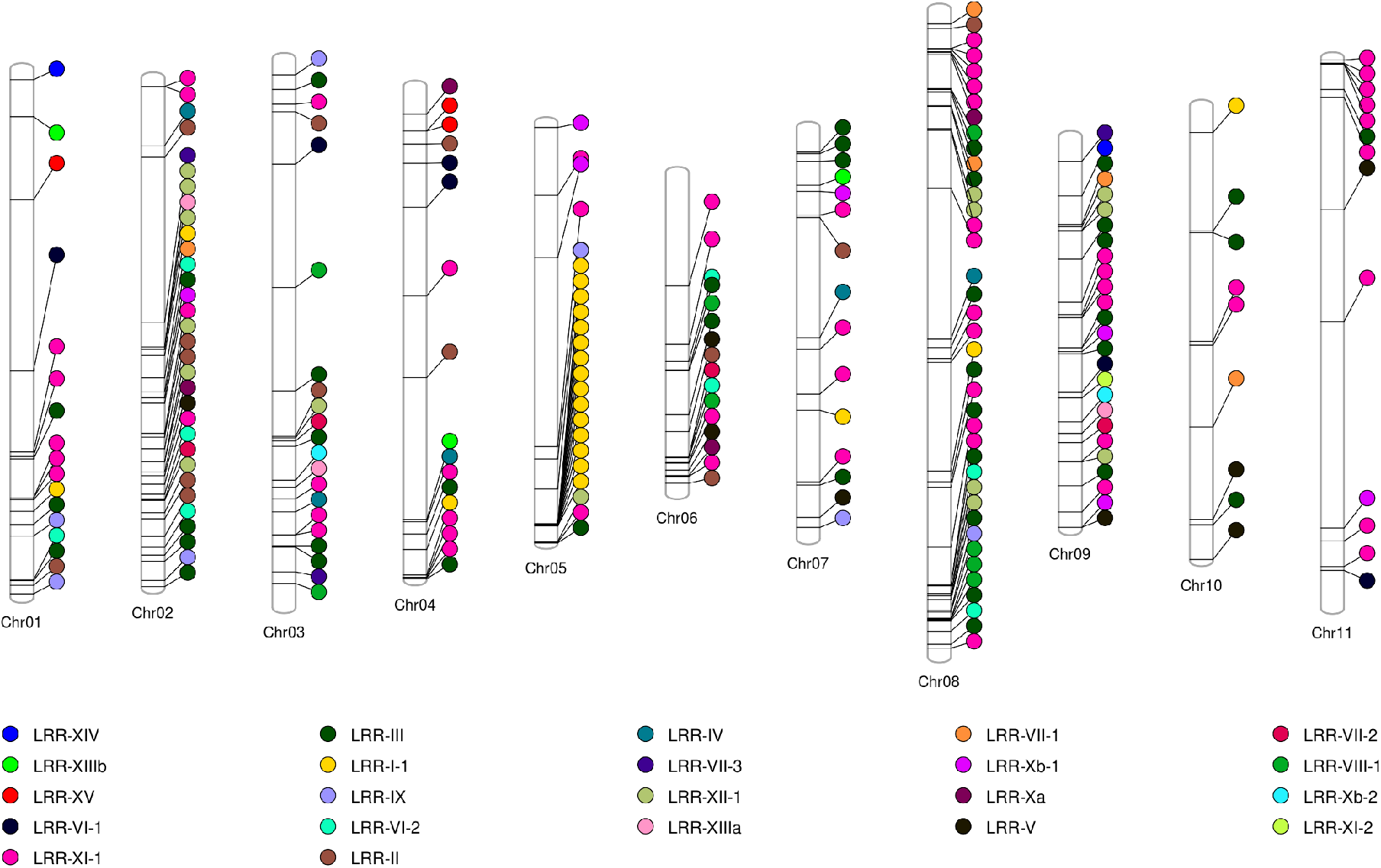
Chromosomal location of the genes encoding LRR-RLKs of *Phaseolus vulgaris* in the your 11 chromosomes, highlighting their link to the subfamily represented by the colors.

**Table 1.**
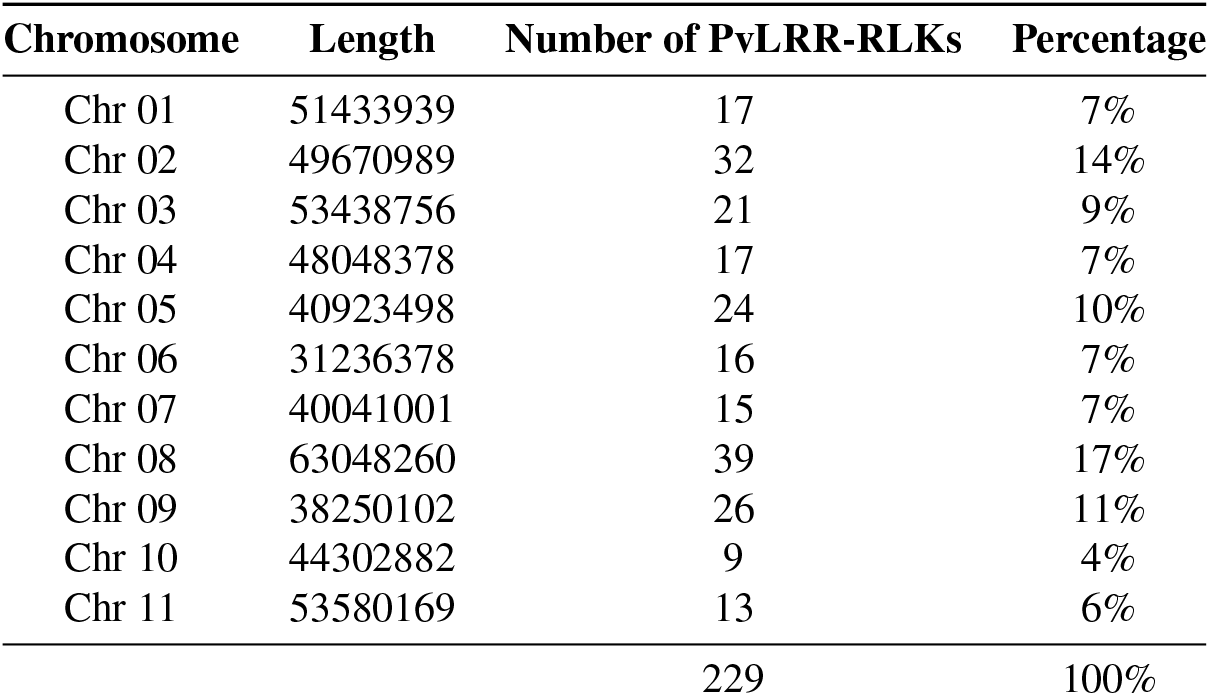
LRR-RLK genes distribution across chromosomes of *Phaseolus vulgaris*.

| Chromosome | Length | Number of PvLRR-RLKs | Percentage |
| --- | --- | --- | --- |
| Chr 01 | 51433939 | 17 | 7% |
| Chr 02 | 49670989 | 32 | 14% |
| Chr 03 | 53438756 | 21 | 9% |
| Chr 04 | 48048378 | 17 | 7% |
| Chr 05 | 40923498 | 24 | 10% |
| Chr 06 | 31236378 | 16 | 7% |
| Chr 07 | 40041001 | 15 | 7% |
| Chr 08 | 63048260 | 39 | 17% |
| Chr 09 | 38250102 | 26 | 11% |
| Chr 10 | 44302882 | 9 | 4% |
| Chr 11 | 53580169 | 13 | 6% |
|  |  | 229 | 100% |

The number of introns in PvLRR-RLKs ranged from 0 to 26 (Supplementary Fig. S4; Supplementary Table S4). We observed 14 genes with no introns in their composition, whereas 100 genes had only one intron, and 24 genes had two introns; showing that the PvLRR-RLKs tend to have few introns, with specific divergences. Proteins Phvul.001G038400, Phvul.004G126100, and Phvul.007G063200 (all the representatives from the subfamily LRR-XIIIa) presented the highest number, with 26 introns. In addition, subfamilies LRR-VII-1, LRR-VII-3, LRR-Xb-1, LRR-XIIIa, LRR-XI-2, LRR-IV, LRR-V, LRR-Xb-2, and LRR-VI -1 showed proteins with the same number of introns in their composition. The other subfamilies showed slight variations in the number of introns. For example, subfamily LRR-I-1 showed the greatest variation of this characteristic among its proteins, with representatives having 8, 11, 13, and 14 introns.

Regarding the promoter region, 115 cis-elements were identified as potential regulators of the expression of the 230 PvLRR-RLKs (Supplementary Fig. S5). Among the typical elements found, the three most frequent were TATA-box (40.17%), CAAT-box (22.86%), and AT∼TATA-box (3.98%). These elements were identified one or more times in all genes. Elements such as Myb (2.65%), Box 4 (2.56%), MYC (2.21%), ERE (1.38%), G-box (1.33%), and ARE (1.32%) were also frequent in the PvLRR-RLK genes. TC-rich repeats, related to defense and stress responses, were identified 83 times in these genes. A diversity of elements related to light stimuli response, such as LAMP motif, L-BOX, ME, TCT motif, AF1 binding site, GT1-1, CMA chs 1a, and 2a^47^ were found in the regions of the studied PvLRR-RLK genes. Furthermore, promoters related to the circadian cycle, low temperature, drought inducibility, and cell cycle are also present in the structure of these genes. These results highlight how complex cis-elements can be, considering that several elements can be present in the same gene.

Bisneta and Gonçalves-Vidigal^51^ collect information present in the literature on loci and QTLs associated with anthracnose resistance and created a map associating the markers located 500 kb upstream and downstream of these regions, based on the common bean reference genome (version 2.1). We used information about marker positions and associated them with the position of the PvLRR-RLKs analyzed in this study (Supplementary Fig. S6). It was possible to observe the presence of markers close to regions where these proteins are concentrated, with emphasis on the telomeric region of the Chr 01 and Chr 05. In Chr 01, the Phvul.001G245300 gene is found between single nucleotide polymorphisms (SNPs), kompetitive allele specific PCR (KASP), and sequence-tagged site (STS) and on Chr 05, there is a SNP marker just below a cluster of genes from the LRR-I-1 subfamily. The distance in base pairs between the RLK-LRR genes of *P. vulgaris* and the markers associated with anthracnose resistance loci are shown in the Supplementary Table S5.

### Duplication Events and Synteny Analysis

When analyzing the duplication events of the LRR-RLK genes throughout the *P. vulgaris* genome (Fig. 4), we observed correspondences between several proteins in more than one chromosome. In total, 178 PvLRR-RLK protein pairs were identified as duplicate segments, corresponding to 77.4% of the total proteins identified in *P. vulgaris*. The collinearity events, Ka/Ks values, and the duplication times were estimated for the PvLRR-RLKs (Supplementary Table S6). Out of the 178 proteins, 175 proteins showed Ka/Ks values lower than one. While the other three (Phvul.005G054300, Phvul.004G037500, and Phvul.001G174500) showed a ratio of more than one. About the duplication time, the Phvul.001G140900 protein was the most recent, having about 32 million years of duplication. On the other hand, Phvul.002G302600 protein has a report of its duplication in approximately 229 million years.

**Figure 4.**
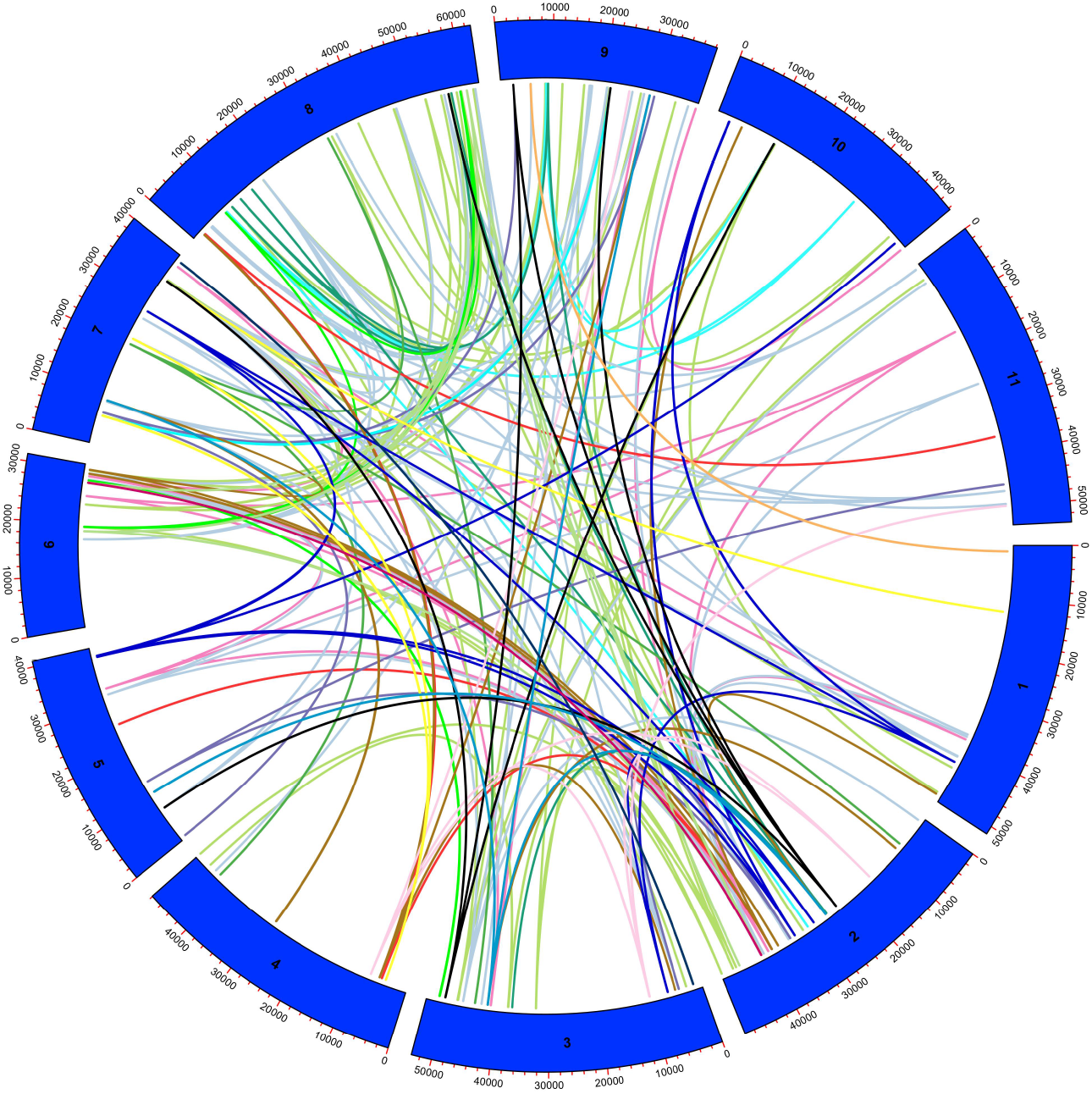
Analysis of duplicated LRR-RLK proteins in *Phaseolus vulgaris*. In the circle, in blue, the chromosomes of *P. vulgaris* are represented by numbers from one to ten. Duplicated proteins are identified by the colored lines inside the circle, indicating their location on the chromosomes.

The evolutionary relationships within the members of these subfamilies were explored by synteny analysis of the PvLRR-RLK genes in *Glycine max*. We found homologies between PKs in all bean and soybean chromosomes (Fig. 5). Out of the 229 PvLRR-RLKs, 172 showed a syntenic relationship with soybean genes, with only 32 having a syntenic relationship with a single gene in soybean, and the majority with more than one gene (Supplementary Table S7). The gene Phvul.003G129400 (Chr 03) exemplifies such a fact, being part of the group of genes that showed a syntenic relationship with five soybean genes, each one located in a different chromosome, e.g., Glyma.07G041200 (Gm07), Glyma.13G111800 (Gm13), Glyma.15G179300 (Gm15), Glyma.16G009900 (Gm16), and Glyma.17G047900 (Gm17).

**Figure 5.**
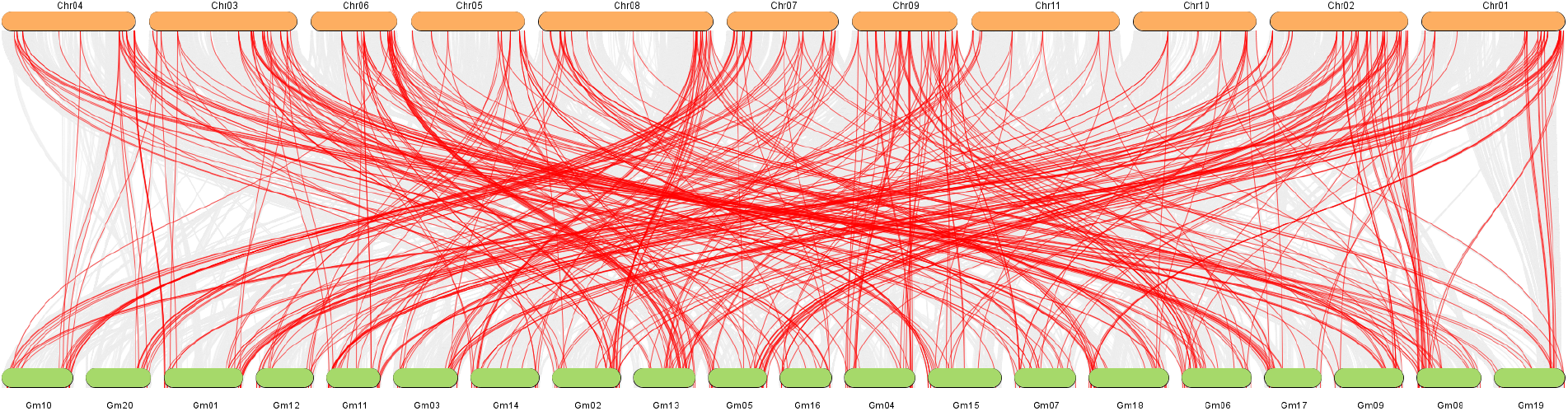
Synteny analysis between the LRR-RLK genes of *Phaseolus vulgaris* and *Glycine max*. The chromosomes of *P. vulgaris* are identified by Chr followed by the chromosome number. The chromosomes of *G. max* are identified by Gm followed by the chromosome number. Thus, Chr 01 corresponds to chromosome 1 of *P. vulgaris*, whereas Gm 01, corresponds to chromosome 1 of *G. max*.

### Study of LRR-RLKs Associated with *P. vulgaris* Resistance to Anthracnose

Transcriptome data of *P. vulgaris* interacting with *C. lindemuthianum* were employed in our study to evaluate the gene expression profile of PvLRR-RLK genes under biotic stress. Padder et al.^53^ used two lines, one resistant (Puebla 152) and other susceptible (Jaguar) to race 73 of *C. lindemuthianum*.

The phred quality values (q value) were greater than 20, being within the quality standard. It was not necessary to clean the sequences, as they were already presented without adapters. We found a total of 948 DEGs in the resistant line at 72 hpi, when compared to the control time 0 hpi. Of the 443 up-regulated DEGs, 2.26% (10 DEGs) corresponds to PvLRR-RLKs, while for down-regulated proteins, of the 495 DEGs, 3.63% (18) are characterized as PvLRR-RLK (Fig. 6A; 6C). Considering the susceptible line, at the time of 72 hpi, compared with the time control at 0 hpi, 4,303 DEGs were found. Of these, 2,549 were up-regulated, 1.84% (47) were PvLRR-RLKs (Fig. 6B). Considering the down-regulated, 1,754 DEGs were found, 1.08% (19) were PvLRR-RLKs (Fig. 6C). Interestingly, it was observed that the 10 up-regulated genes in the resistant line, at 72 hpi, were also up-regulated in the susceptible line, which in turn had 37 unique up-regulated genes, at the same time (Figure 6C).

**Figure 6.**
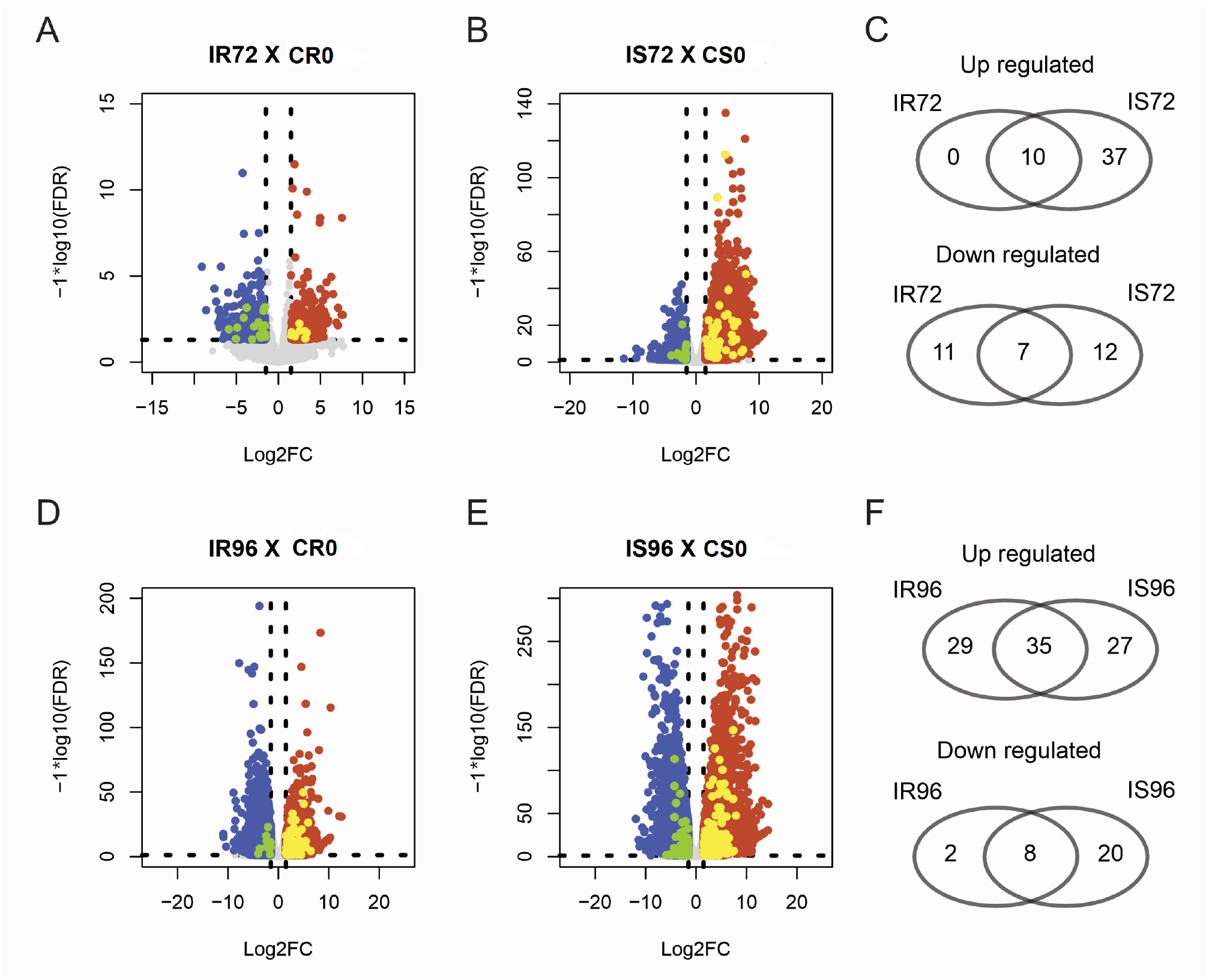
Analysis of LRR-RLK genes of *Phaseolus vulgaris* involved in the response of resistant (Puebla 152) and susceptible (Jaguar) lines to race 73 of *Colletotrichum lindemuthianum* at 72 and 96 hours after inoculation (hpi) compared with the time control 0 hpi. IR72 represents the resistant line inoculated at 72 hpi. IR96 represents the resistant line inoculated at 96 hpi. CRO represents the control resistant line inoculated at 0 hpi. IS72 represents the susceptible line inoculated at 72 hpi. 1S96 represents the susceptible line inoculated at 96 hpi. CS0 represents the control susceptible line at 0 hpi.

At 96 hpi, 4,587 genes were differentially expressed between the resistant line and the control time 0 hpi. Of the 4,587 DEGs, 2,659 were up-regulated in the resistant line, and 2.40% (64) were PvLRR-RLKs. Out of the 1,928 down-regulated in the resistant line, 0.52% (10) were PvLRR-RLKs (Fig. 6D, 6F). Comparing the 96 hpi susceptible line with the 0 hpi control time, there were 4,275 up-regulated DEGs and 3.327 down-regulated DEGs in total, being of those 1.45% (62) and 0.84% (28) PvLRR-RLKs up-regulated and down-regulated, respectively (Fig. 6E, 6F). The relationship of up-regulated and down-regulated proteins at each time in the two lines is shown in Supplementary Tables S8, S9.

The profile of the fold-change log2 values (based on the TPM values) of the PvLRR-RLKs showed a significant difference in at least one of the interaction times (72 or 96 hpi) in resistant and susceptible plants, which were highlighted in a heatmap cluster analysis. This filter excluded 75 analyzed genes from the analysis. The other genes were separated into five groups (A to E) according to the expression pattern presented at the different analyzed times. In group A, for example, there are genes that, in general, were down-regulated or without expression at 72 hours in the resistant line and that, at 96 hours, started to be up-regulated. Groups B and C stand out for down-regulated genes in the two analyzed times, in both lines. While in groups D and E, there are genes up-regulated, at 72 and 96 hpi, in resistant and susceptible lines. Of D and E groups, the genes grouped in D stand out for presenting higher fold-change log2 values, mainly in susceptible line (Fig. 7).

**Figure 7.**
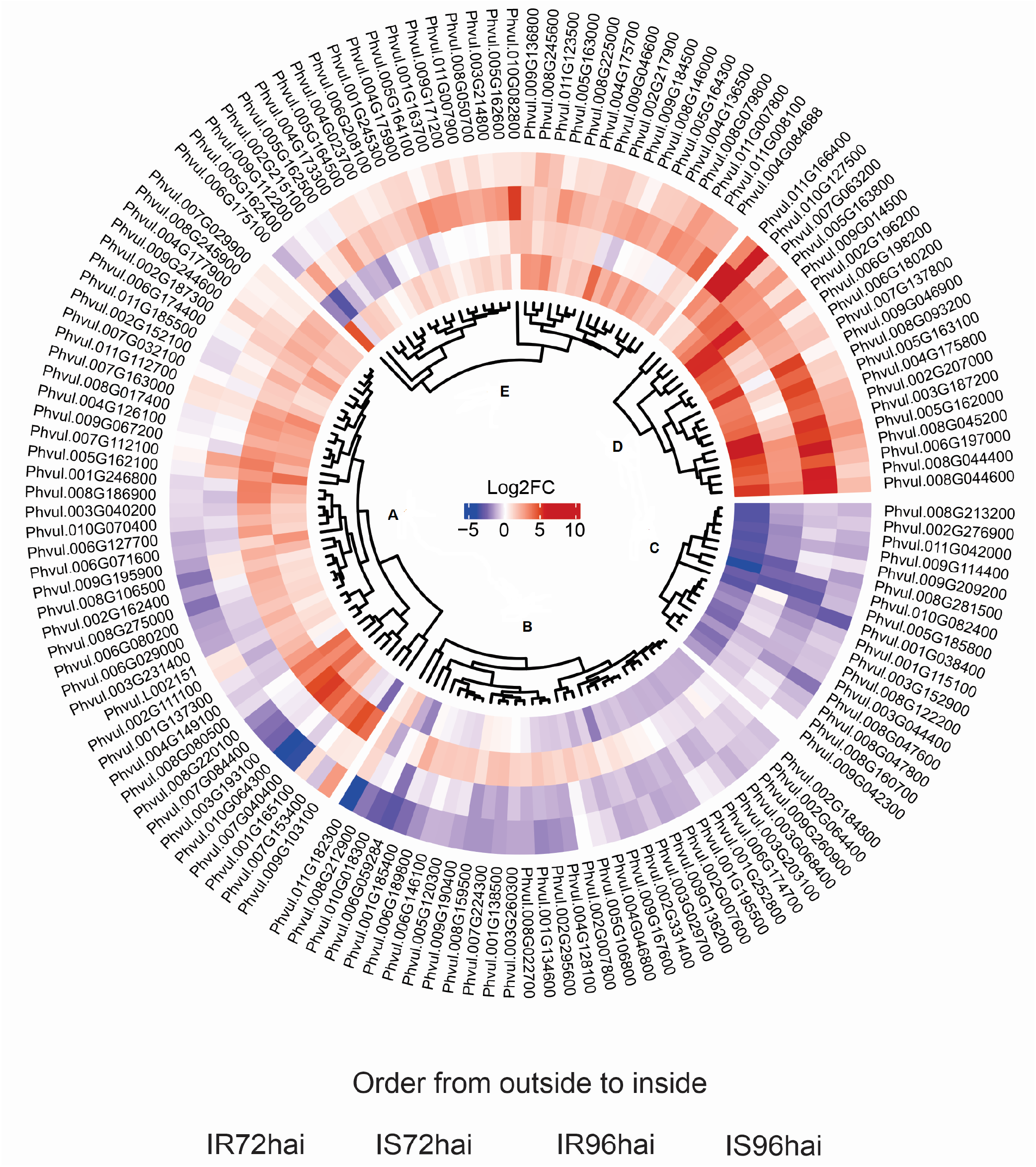
Heatmap analysis of differentially expressed LRR-RLK genes of *Phaseolus vulgaris* in resistant (Puebla 152) and susceptible (Jaguar) lines inoculated with race 73 of *Colletotrichum lindemuthianum* at 72 and 96 after inoculation (hpi).

## Discussion

LRR-RLK proteins promote key membrane-anchored receptors responsible for recognizing external factors, transducing apoplastic signals into symplasts, and triggering responses to different stimuli^60^. In our study, we identified and characterized 230 PvLRR-RLKs (Supplementary Table S2). The approximate numbers had already been identified in species such as *A. thaliana* (239)^61^ and *Oryza sativa* (309)^62^. The PvLRR-RLK proteins identified vary in physicochemical properties such as molecular weight, isoelectric point, and gene size (Supplementary Table S3). These features may result from the different functions developed in their own microenvironments. In general, LRR-RLKs constitute one of the largest gene families, from the class of KD superfamily, in the animal and plant kingdoms^63^, which can be related to their critical role in different fundamental biological processes, e.g., signal transduction, innate cell immunity, growth and development, cell differentiation and patterning, nodulation, and self-incompatibility mechanisms^64, 65^.

The identified proteins were classified into 15 subfamilies, some of which were subdivided (Supplementary Table S3). The classification pattern obtained was similar to *A. thaliana, B. rapa*, and *O. sativa*^24^. The phylogenetic tree was estimated in order to confirm the classification obtained for the LRR-RLK proteins. There was a coincidence of both classifications for all subfamilies, except LRR-XI-1, which were divided into two subclades in the tree. However, all other subclades were well-supported, with statistical confidence values above 0.70 (Fig. 1). Such values suggest reliability in the data obtained^16^. The *C. reinhardtii* LRR-RLK protein was used as an outgroup. Evolutionary analysis of the RLK superfamily indicates the existence of this family before the divergence of land plants and algae^40^, and therefore it was used to root the tree in this study.

Conserved domain and motif analysis was performed to structurally characterize these proteins. The presence and organization of the domains and motifs was consistent with the phylogenetics analysis, which corroborates the division into subfamilies (Supplementary Fig. S1; S2). A similar pattern was obtained for proteins of this same family in paper mulberry, *Cucumis sativus*^48,66^, and other gene families, e.g., the WRKY family in wheat^67^. As for domain analysis, the malectin domain was found in proteins classified as subfamily LRR-I-1 (Supplementary Fig. S1). Current research in plant models suggests that proteins with malectin/ malectin-like domains function as multiple wall sensors involved in processes that depend on or affect the cell wall in various ways, e.g., growth control reproduction and multiple stresses^68^. This domain was also found in 13 LRR-RLK proteins in cotton^60^ and Arabidopsis^69^, which were grouped into the same LRR-1 subfamily. Hu et al.^70^ reported that the RLK-V malectin/leucine-rich repeat receptor protein kinase gene from *Haynaldia villosa* acts as a PRR to up-regulate resistance to powdery mildew caused by *Blumeria graminis* f. sp. *tritici* (Bgt) in wheat. When analyzing the conserved motifs present in the KD and LRR domains, we found 15 (Supplementary Fig. S2). Similar numbers were found in paper mulberry, with 13 motifs^48^, and in cucumber, with 12 motifs^66^. In some subfamilies, one or two proteins showed a different motif 10 from the others. This motif is associated with the LRR 4 domain. Other motifs were also associated with this same domain, e.g., motif 7 and motif 12. Due to the diverse functional roles of LRR-RLK proteins, these proteins have specialized domains and motifs for functional specializations^69^.

One way to gain insights into possible functions of a protein is to analyze its subcellular localization, considering that most of the biological activities performed by proteins are closely related to where they reside in the cell^71^. Upon further analysis at the cellular level, more than 63% of the PvLRR-RLKs were related to the plasma membrane (Supplementary Fig. S3), corroborating their main functions of recognizing environmental stimuli and internal signals^16^. Except for subfamilies LRR-I-1, LRR-XIV, LRR-VII-3, and LRR-XIII-a, all others showed proteins grouped in different locations, showing that this parameter is not a determining factor in protein grouping. Differences in the extracellular domains and associated structure resulted in the functional specialization of these proteins and, consequently, in their cellular localization^60, 72^. For example, only the Phvul.009G043600 protein (subfamily LRR-VIII-1) showed cytoplasmic localization. Coincidentally, it is the only protein in the subfamily that shows a different KD domain, the Kinase-like superfamily, and a tripled LRR 8 domain.

Starting with the characterization of the members of the LRR-RLK family in *P. vulgaris* at the genomic level, we analyzed the gene structure, based on the exon/intron organization and analysis of cis elements. The PvRLK-LRRs genes had from zero to 26 introns in their structure (Supplementary Fig. S4). The same number range was also found for this gene family in *Populus* and *Arabidopsis*^23^. There was no direct relationship between the number of introns found in genes grouped into the same subfamily. However, the variation of this parameter in the same group was small. This pattern of slight variations in gene structure was also found for this same group of proteins in *Populus* and *Arabidopsis*^23^.

For analysis of the promoter regions, 1.5 kb upstream was used. Most cis elements are composed of five to 20 nucleotides and are located in the proximal promoter, a region that encompasses several hundred nucleotides upstream from the transcription start site^28,73^. The TATABOX and CAAT-BOX elements were the most frequently found (Supplementary Fig. S5). Both elements are important sequences of promoter regions, the first being a nucleotide sequence containing approximately 30 nucleotides before the transcription start site^74^, whereas the second represents a common cis-action element in promoter and enhancer regions, containing about 40 to 100 nucleotides above the transcriptional start site^75^. MYC elements frequent in the promoter regions of PvLRR-RLK genes function primarily in growth and development, as well as in response to stress^76^. In addition, elements related to hormone responses, e.g G-box, ABRE, and ERE, were also identified in these genes. These cis-elements are potentially responsive to abscisic acid, ethylene, and methyl jasmonate. They were also found in large numbers in cotton LRR-RLK genes^47^.

The LRR-RLK genes are distributed in all chromosomes of *P. vulgaris*, especially in their telomeric regions (Fig. 3). When two or more genes of the same family are located in a chromosomal region of about 200 kb, these are defined as clusters, whereas genes that share an identity of more than 70% in this situation are considered to result from tandem duplication^38,77^. Gene distribution is something to be highlighted, considering its informative power for understanding linkage groups and the possibilities of recombination. As for the location of PvLRR-RLKs on chromosomes, Schmutz et al.^78^ observed genes related to disease resistance in clusters at the end of Chr 04, Chr 10, and Chr 11 in *P. vulgaris*. Of these, in our analyses, Chr 04 presented the highest number of PvLRR-RLKs in these regions. In addition, kinases at the *Co*-4 of *P. vulgaris* locus have been reported to provide PTI^53^. Bisneta and Gonçalves-Vidigal^51^ also found that most genes encoding kinase proteins related to anthracnose resistance were located in clusters at the end of the chromosomes, e.g., cluster 1.1 on chr 01. The Phvul.001G245300 gene, subfamily LRR-VI-2, was located among different types of markers (Supplementary Figure S6; Supplementary Table S5), and can be considered, due to its location close to the *Co*Pv01cdrk locus, a candidate for anthracnose resistance^51^.

To analyze the evolutionary aspect of PvLRR-RLK genes, we performed duplication and synteny analysis. More than 75% of the genes identified correspond to duplicated segments, suggesting that the formation of some PvLRR-RLK genes may have arisen from duplication events (Fig. 4; Supplementary Table S6). As in *Populus*, approximately 82% of the LRR-RLKs genes are also located in duplicated regions^23^. The Ka/Ks ratio was performed, and of the 178 proteins, only three (Phvul.005G054300, Phvul.004G037500, and Phvul.001G174500) showed values higher than one for this parameter, indicating positive selection. In contrast, all the other 175 proteins showed values lower than one, suggesting that most LRR-RLKs genes of *P. vulgaris* may have passed through stabilizing selection during the evolution process. A similar pattern was observed for the potato bZIP gene family^38^.

Considering the synteny analysis, common bean and soybean are the two most economically important members of the Phaseoleae legumes, soybean for its many human and animal usages, and common bean as an important nutritional crop for many economically poorer countries^79^. LRR-RLK genes showed a high homology index (Fig. 5; Supplementary Table S7), highlighting the close relationship between the two species. The relationship indicates that these genes may have existed before the differentiation of the two species and have maintained a collinear relationship since then^38^. Anthracnose is also an economically important disease for soybean, and can strongly affect its production, under suitable conditions for the development of the pathogen^80^. Therefore, extrapolating the understanding of this protein subfamily to soybean, through the analysis of orthologous genes, constitutes an important tool in obtaining resistant cultivars, one of the main control methods used.

The expression analysis of PvLRR-RLK genes using data from Padder et al.^53^ was carried out to verify the expression profile of this gene family, which is often related to stress responses such as diseases in a pathosystem^28^. The strong expression profile of some DEGs draws attention to the image (Fig. 7). For example, the genes Phvul.011G166400 and Phvul.008G045200 were strongly expressed in the susceptible line at 72 hours. The first is associated with a nuclear protein located on Chr 11 and belongs to the subfamily LRR-XII, whereas the second was grouped in the LRR-XI-1 subfamily, representing a plasma membrane protein located on Chr 08. Possibly, these genes could be somehow associated with the onset of symptoms.

In the present investigation, many PvLRR-RLK genes showed an interesting down-regulated or not expressed gene pattern in the resistant line at 72 hpi, with their expression increasing with the time of pathogen infection (Fig. 7A). Of these genes, we highlight some that were located in Chr 01, Phvul.001G246800 (subfamily LRR-III) and Phvul.001G165100 (LRR-XI-1), the first being located between the SNP (SS83) and STS (TGA1.1570) markers, and the second, close to the SNP marker, PvM15. These marks are located near resistance loci^51^. Both genes represent a plasma membrane protein. Even in Chr 01, the Phvul.001G245300 gene (LRR-XI-2) was up-regulated in the resistant line, at 72 hpi, when compared to the susceptible line, and its expression decreased over time, at 96 hpi (Fig. 7E). This gene is a cytoplasmic protein. With the exception of the Phvul.001G165100 gene, the others had already been cited as resistance candidates^51^, however, due to its proximity to a marker associated with anthracnose, and its expression pattern, we believe that it represents a putative element in the common bean resistance response to the disease. In Chr 04, the genes Phvul.004G149100 (LRR-III/ membrane plasmatic protein) and Phvul.004G128100 (LRR-IV/ extracellular protein) were located close to the SNP marker NSSR65, and showed a higher level of expression in the 96 hpi resistant line when compared to the susceptible line, at the same time. They were present in the genomic region of the *Co*-1, *Co*-x, and *Co*-4 genes. These chromosomes have already been reported as important resistance clusters^51^.

At the end of Chr 05, the presence of 15 genes in tandem from the same subfamily, LRR-I-1, caught our attention. When analyzing the expression pattern of these genes, we found 11 DEGs. Of the 11 DEGS, six (Phvul.005G162400, Phvul.005G162500, Phvul.005G162600, Phvul.005G163000, Phvul.005G164300, and Phvul.005G164500) showed the same pattern, being downregulated at both 72 and 96 hpi, in both lines (Fig. 7, group E). The genes Phvul.005G162000, Phvul.005G163100, and Phvul.005G163800 were up-regulated in all treatments, however, the first two showed higher expression levels at times of 72 and 96 hpi in the susceptible line, and the last one, in the two times of the resistant line (Fig. 7D). These genes are close to the SSR marker PvM07, previously associated with anthracnose resistance locus^51^. Still considering this subfamily, in addition to Chr 05, there are DEGs distributed in Chr 02, 07, 08 and 10. In Chr 07, the gene is close to the scaffold marker 00098_217812 (SNP), which is up-regulated in resistant and susceptible lines at 96 hpi, when compared with the same treatments at 72 hpi. In Chr 10, the gene is close to the SNP marker, ss715648754, and was only expressed in the treatments of the line resistant to 72 hpi and susceptible to 96 hpi, which was down-regulated (Supplementary Fig. S6). All the genes of this family present the malectin domain and correspond to membrane plasmatic proteins. Despite the proximity to some markers, so far, no gene from this subfamily has been associated as a candidate for anthracnose resistance.

One of the most well-reported examples in the literature is the protein resulting from the expression of the FLS2 gene, identified as Phvul.002G196200 and grouped in subfamily LRR-XII-1 (Fig. 1). This protein contains one intron and is located on Chr 02 of the *P. vulgaris* genome. According to the heatmap, the FLS2 gene is more expressed at 96 hpi in both lines but stands out in the susceptible line. Identified in *Solanum lycopersicon, Brassica, Arabidopsis*, and *Oryza sativa*^22,25,27,81^, the FLS2 gene is involved in the perception of the bacterial elicitor FLAGELIN and acts as a PRR in the initial plant defense response^82^. The expression pattern found by Padder et al.^53^ shows that the gene is upregulated in all treatments, standing out, however, in resistant and susceptible lines at 96 hpi. Oblessuc et al.^83^ observed the downregulation of this gene at 65 hpi with race 73 of *C. lindemuthianum*, indicating a progressive curve of this gene over time. On the contrary, Silva et al.^20^ evaluated the expression of genes related to the resistance of common bean to race 65 of *C. lindemuthianum* and obtained a positive regulation of this gene with the BRS Esplendor line, characterized by its resistance to the studied fungus at 72 hpi. In contrast, there was no differential expression between the inoculation and control treatments for the pathosystem involving the susceptible line. Although they are the same crop and the disease, the pathogenic variability between and within races is possibly a determining factor for developing a specific response, with the same genes acting differently.

In our study, we performed a full characterization of 230 PvRLK-LRR proteins, supplying different insights into their evolutionary history. In addition, we integrated our findings with the expression profile of PvLRR-RLK genes in response to the infection with *C. lindemuthianum*, investigating the proximity of such genes with markers associated with resistance loci. This information is important, as it allows insights into the role of these genes in the common bean-anthracnose pathosystem. These candidate genes may be useful for further studies to validate their functions in ANT response and to understand how they interact within metabolic pathways.

## Acknowledgements

The authors gratefully acknowledge the Fundação de Amparo à Pesquisa do Estado de Minas Gerais (FAPEMIG), the Coordenação de Aperfeiçoamento do Pessoal de Nível Superior (CAPES), and the Conselho Nacional de Desenvolvimento Científico e Tecnológico (CNPq) for for financial support, and the Fundação de Amparo à Pesquisa do Estado de São Paulo (FAPESP) for a Ph.D. fellowship to AA (2019/03232-6)

## Author contribution statement

Conceived and designed the analysis: CMSD, WAP, AA. Performed the analysis: CMSD, WAP, AA, EMAS. Wrote the original draft preparation: CMSD. Wrote review and editing: CMSD, WAP, AA, EMAS.

## Data Availability

All data generated or analyzed during this study are included in this published article and its supplementary information files.

## Supplementary Tables

**Supplementary Table S1**. Kinase and LRR domains annotation of 230 proteins.

**Supplementary Table S2**. Subfamily classification of bean 230 LRR-RLKs.

**Supplementary Table S3**. Bean 230 LRR-RLKs compositional analyses.

**Supplementary Table S4**. Bean 230 LRR-RLKs subcellular localizations.

**Supplementary Table S5**. Position of LRR-RLK genes and molecular markers associated with resistance loci in *Phaseolus vulgaris* chromosomes.

**Supplementary Table S6**. Collinearity events and Ka/Ks values of bean LRR-RLKs.

**Supplementary Table S7**. Synteny between *Phaseolus Vulgaris* and *Glycine max* genes.

**Supplementary Table S8**. Bean RLK-LRR genes differentially expressed at 72 hours after inoculation in strains Puebla 152 (resistant) and Jaguar (susceptible) with race 73 of *Colletotrichum lindemuthianum*.

**Supplementary Table S9**. Bean RLK-LRR genes differentially expressed at 96 hours after inoculation in strains Puebla 152 (resistant) and Jaguar (susceptible) with race 73 of *Colletotrichum lindemuthianum*.

## Supplementary Figures

**Supplementary Fig. 1**. Phylogenetic and functional domain analyses of LRR-RLK proteins from *Phaseolus vulgaris*. In the phylogenetic tree, the different colors represent distinct protein classifications into subfamilies obtained by HMMER. The different domains found are also identified by different colors.

**Supplementary Fig. 2**. Phylogenetic and conserved motif analyses of LRR-RLK proteins from *Phaseolus vulgaris*. In the phylogenetic tree, the different colors represent distinct protein classifications into subfamilies obtained by HMMER. The different motifs found are also identified by different colors.

**Supplementary Fig. 3**. Phylogenetic analysis and subcellular localization heatmap of LRR-RLK proteins from *Phaseolus vulgaris*. In the phylogenetic tree, the different colors represent distinct protein classifications into subfamilies obtained by HMMER. The most likely subcellular localization is indicated in parentheses next to the protein identification.

**Supplementary Fig. 4**. Phylogenetic and gene organization (exon and intro compositions) analyses of LRR-RLK genes from *Phaseolus vulgaris*. In the phylogenetic tree, the different colors represent distinct protein classifications into subfamilies obtained by HMMER.

**Supplementary Fig. 5**. Phylogenetic and cis-element analyses of LRR-RLK genes from *Phaseolus vulgaris*. In the phylogenetic tree, the different colors represent distinct protein classifications into subfamilies obtained by HMMER. The different cis-elements found are also represented by different colors.

**Supplementary Fig. 6**. Chromosomal colocalization of *Phaseolus vulgaris* LRR-RLK genes and markers located 500 kb upstream and downstream to anthracnose resistance loci and quantitative trait loci (QTLs).

