## Supplementary figures and images for "Genome-wide analysis and characterization of the LRR-RLK gene family provides insights into anthracnose resistance in common bean"

### S1_domain.tiff

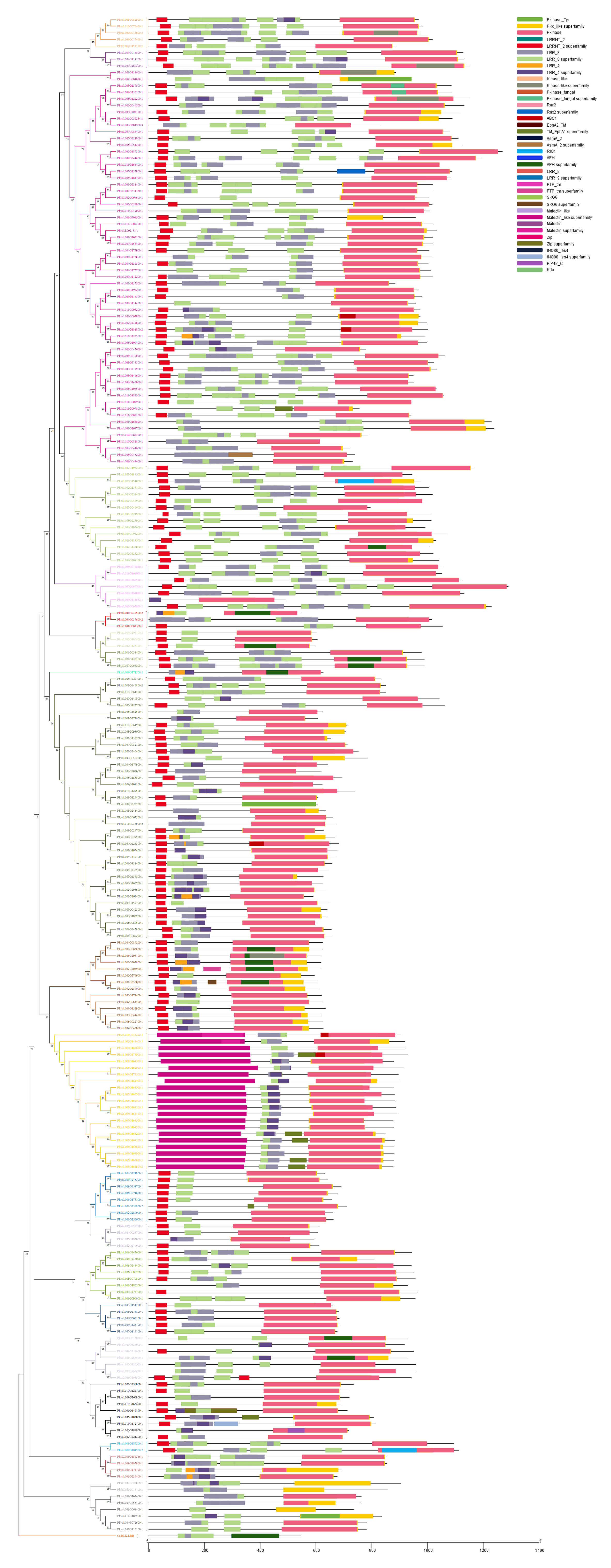

### S2_motifs.tiff

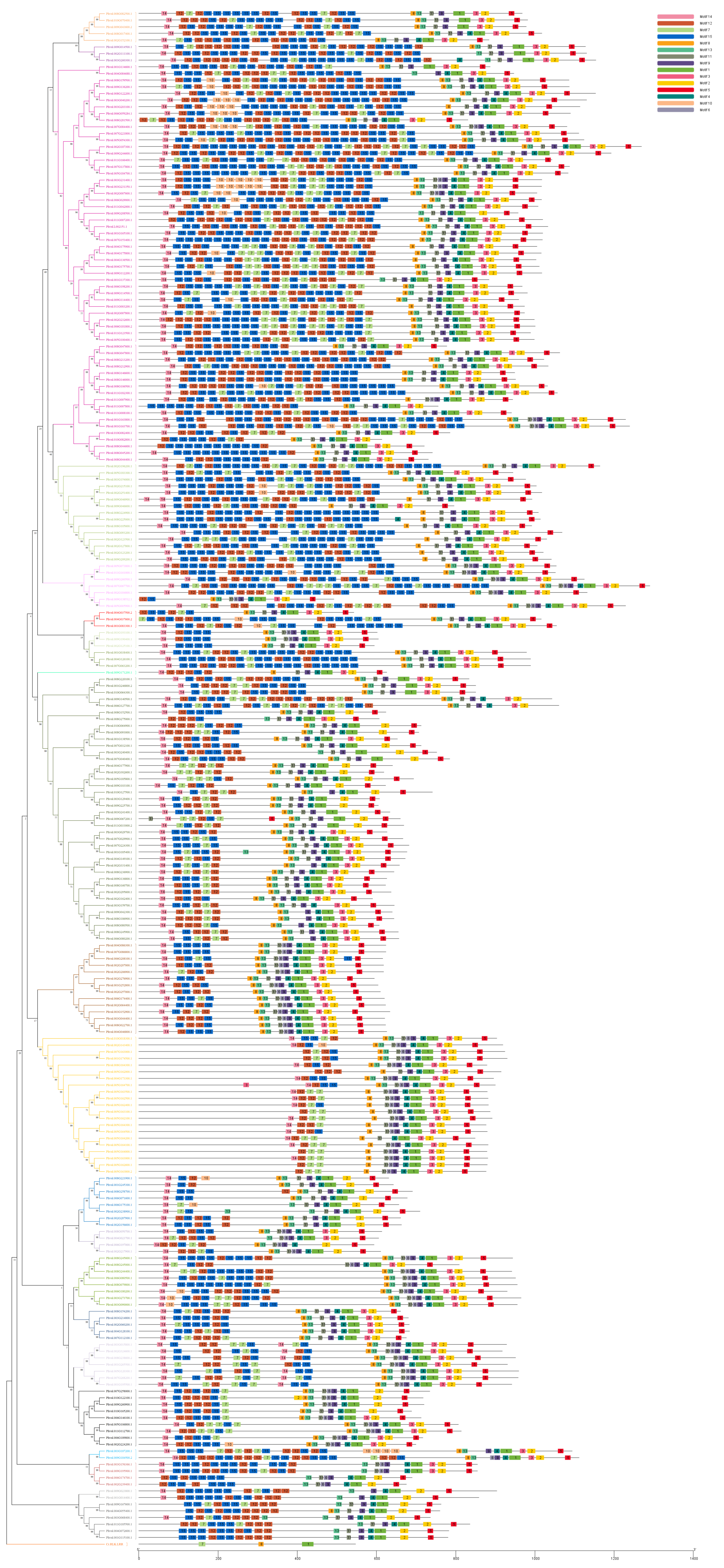

### S3_celular_localization.tiff

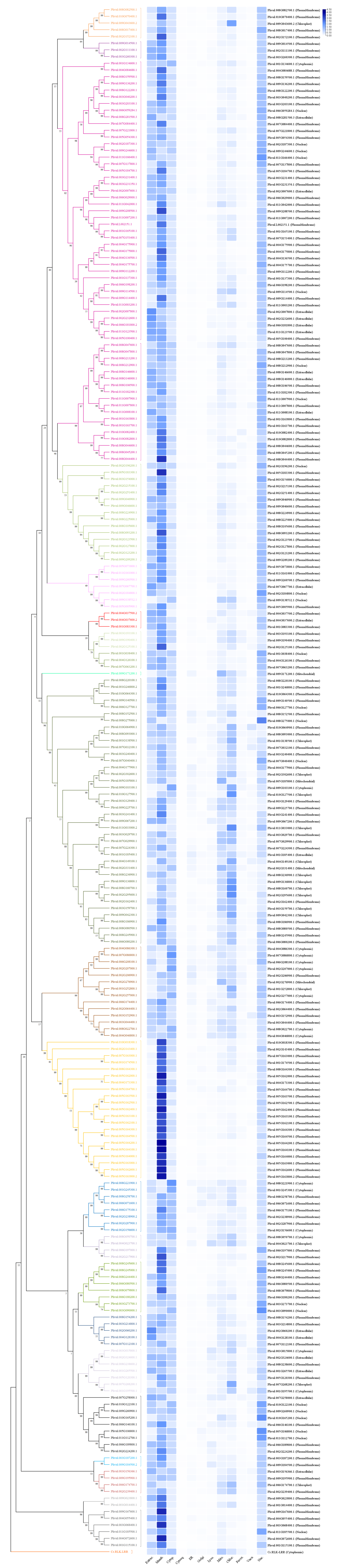

### S4_gene_structure.tiff

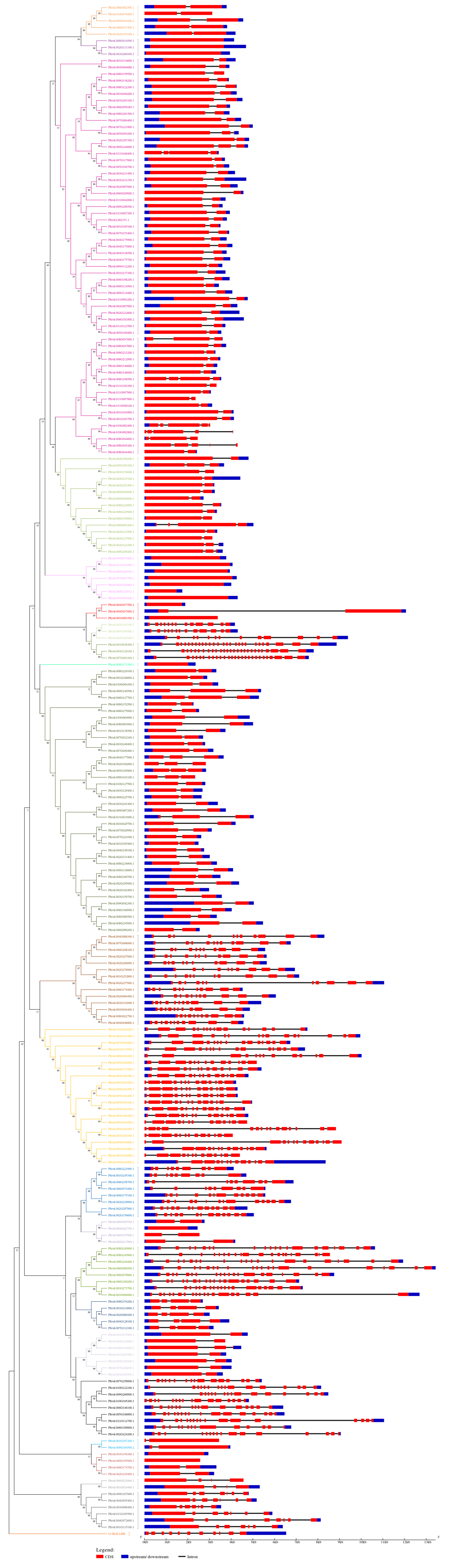

### S6_markers.tiff

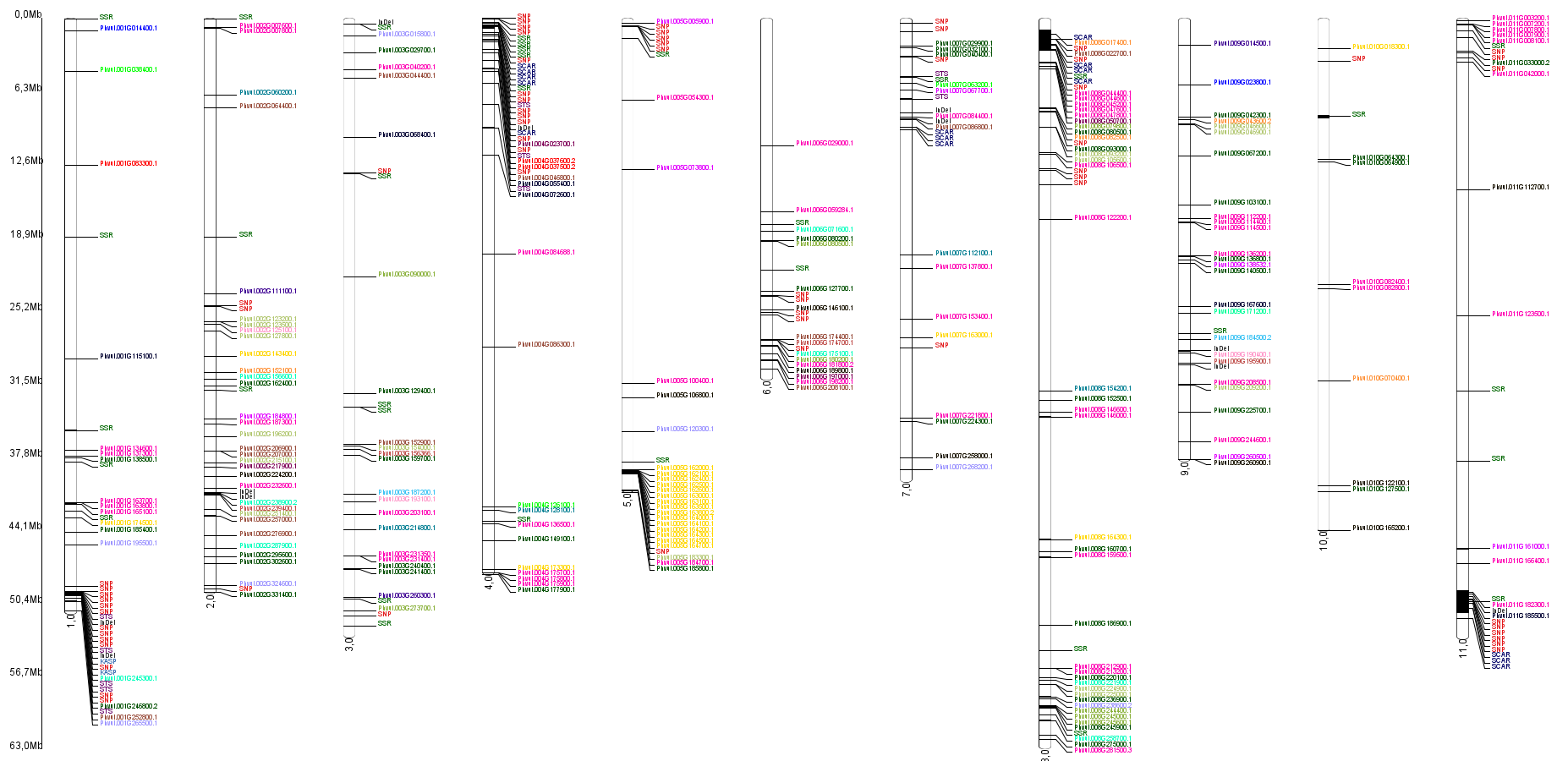
